# Low-salinity medium for large-scale biomass production of the marine purple photosynthetic bacterium *Rhodovulum sulfidophilum*

**DOI:** 10.1101/2025.03.14.643199

**Authors:** Shamitha Rao Morey-Yagi, Dao Duy Hanh, Miki Suzuki, Shota Kato, Geoffrey Liou, Yuki Kuroishikawa, Ayaka Yamaguchi, Hiromasa Morishita, Masaki Odahara, Keiji Numata

## Abstract

Marine purple non-sulfur bacteria such as *Rhodovulum sulfidophilum* are versatile due to their diverse applications in bioremediation, biotechnological production of useful materials, industrial production of value-added compounds, and agricultural fertilizers. Our previous study demonstrated the potential of its lysed and dried biomass as a nitrogen fertilizer for plant production. However, large-scale fertilizer production requires scaling up the culture to larger volumes, which is not cost-effective with currently available options for growth media. In this study, we tested a seawater-based, cost-effective alternative to the commonly used nutrient-rich culture medium for the growth of this bacterium. We found that reducing salinity from 3% to 1.2% had no adverse effects on its heterotrophic growth, dry cell yield, nitrogen content, and total amino acid composition. The nitrogen content and the weight percent of free lysine, aspartic acid, and glutamate increased in the biomass obtained from cultures grown at 1.2% salinity. Under autotrophic conditions, decreasing salinity to 1.2% did not affect cell growth, final dry cell yields, and total carbon assimilation, but N assimilation tended to be higher. Reducing salinity to 1.2% proved to be cost-effective and feasible for the cultivation of *R. sulfidophilum* without increasing the risk of contamination, providing a viable alternative for its large-scale cultivation and application as a plant nitrogen fertilizer.

## Introduction

Marine purple non-sulfur bacteria (PNSB) are photomixotrophs capable of using both CO_2_ (autotrophy) and organic carbon (C) compounds (heterotrophy) to meet their C needs. Many PNSB species can also acquire nitrogen (N) by fixing N_2_ using endogenous nitrogenases (1,2). Additionally, they can produce value-added compounds such as carotenoids (3,4), vitamins (4,5), and polyhydroxyalkanoates (6–10). These bacteria are reported to tolerate a wide range of salinity (4,11,12) and are predominantly grown in culture medium with 2% – 3% NaCl (7,9,10,12–15). The use of natural seawater for their culture has been previously proposed for the sustainable, cost-effective production of these value-added compounds with a low risk of biological contamination (13,16,17). However, this has not been tested yet.

Our earlier studies demonstrated that a gram-negative marine PNSB, *Rhodovulum sulfidophilum* (ATCC 35886), can synthesize polyhydroxyalkanoates (PHA) under photoautotrophic (7–10,17) and heterotrophic (13,15) conditions. It can also be used as a host for the heterologous expression of spider silk (14). Additionally, we have reported that its lysed and dried biomass can be used as a sustainable N fertilizer for plant cultivation (18), allowing biomass waste from the extraction of value-added compounds to be repurposed as plant fertilizer. The high N content of ∼11% and protein content of ∼69% enable its use as a N fertilizer and a proposed amino acid supplement respectively (18). However, large-scale cultivation of this bacterium in a nutrient-rich medium towards this end is not without significant costs. Therefore, in this study, we evaluated the growth, nutrient (N, P, K), and amino acid profile of *R. sulfidophilum* in a low-salinity medium, intending to improve biomass yield and reduce large-scale culture costs, without additional risks of contamination.

## Results

### An artificial seawater-based culture medium is cost-effective for the heterotrophic cultivation of *R. sulfidophilum*

We compared the cell yield and culture media costs of culturing *R. sulfidophilum* in three different growth media namely, commercially available marine broth (MB), a natural seawater (NSW) based and an artificial seawater (ASW) based medium both supplemented with 0.1% yeast extract and 0.5% peptone in a 10 L scale. The fresh cell yield in MB was significantly higher than the seawater-based media, but their dry cell yields were similar (Table 1), indicating that the seawater-based media can replace MB. Compared to the marine broth, the use of NSW- and ASW-based culture media showed a 90% and 91% reduction, respectively, in culture medium costs (Table 1). Culture in the ASW-based medium decreased costs by an additional 9% than when cultured in the NSW-based culture medium (Table 1). Hence, we used the ASW-based culture medium for further salinity tests.

**Table 1.** Cell yield and cost analysis of various growth media for large-scale cultivation of *Rhodovulum sulfidophilum*.

|  | Fresh cell yield<br>(g L<sup>-1</sup>) | Dry cell yield<br>(g L<sup>-1</sup>) | Cost for 1 L<br>(yen) | Dry cell yield<br>relative to MB | Cost relative<br>to MB | Cost relative<br>to NSW |
| --- | --- | --- | --- | --- | --- | --- |
| MB (76448, Millipore) | 6.99 ± 0.76<sup>a</sup> | 0.96 ± 0.10 | 1541 |  |  |  |
| NSW + 0.1% YE + 0.5% Peptone | 4.65 ± 1.10<sup>b</sup> | 1.03 ± 0.23 | 148 | 7% | -90% |  |
| ASW + 0.1% YE + 0.5% Peptone | 5.03 ± 0.68<sup>b</sup> | 0.88 ± 0.10 | 134 | -8% | -91% | -9% |

Data represent means ± SEM from 4 independent 10 L cultures, or batches (n=4). Different letters indicate the significance of differences observed with Marine Broth (MB) as evaluated by a two-way ANOVA (Tukey's test).

### Lower salinity has no negative effects on the heterotrophic growth of *R. sulfidophilum* in ASW-based media

*R. sulfidophilum* was cultured in 15 mL of ASW supplemented with 0.1% yeast extract and 0.5% peptone (Fig. 1a and 1b) or 1 mM sodium acetate and 2.5 mM sodium thiosulfate (Fig. 1c) at decreasing concentrations of ASW i.e. 100%, 90%, 80%, 70%, 60%, and 50% which correspond to 3%, 2.7%, 2.4%, 2.1%, 1.8%, and 1.5% salinities respectively. The optical density of cultures at 660 nm in 60% ASW and 50% ASW was higher than in 100% ASW at 72, 128, and 177 hours and 128 and 177 hours of culture respectively (Fig. 1a). But there were no significant effects of lower salinities (down to 1.5%) on cell dry weight (Fig. 1b). Similar results were also obtained for cultures supplemented with 1 mM sodium acetate and 2.5 mM sodium thiosulfate (Fig. 1c), wherein a decrease in salinity had no negative impact on cell dry weight.

**Fig. 1.**
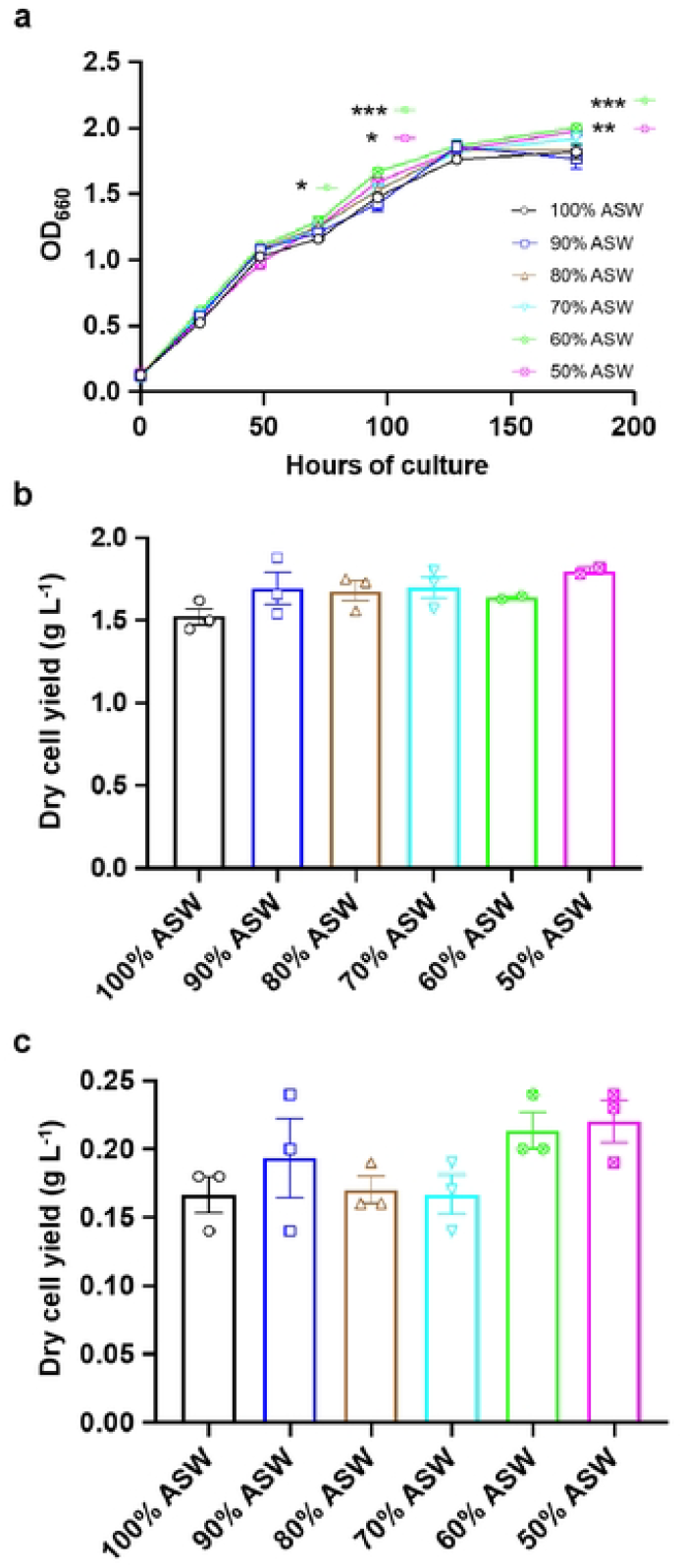
Heterotrophic cell growth and dry cell yield of *Rhodovulum sulfidophilum* in artificial seawater (ASW) based growth medium of different salinities, wherein 100% corresponds to 3% salinity. **a**, Growth curve and **b**, dry cell yield (g L^-1^) of *R. sulfidophilum* in ASW supplemented with 0.1% yeast extract and 0.5% peptone, and **c**, dry cell yield (g L^-1^) of *R. sulfidophilum* in ASW supplemented with 1 mM sodium acetate and 2.5 mM sodium thiosulfate in decreasing concentrations of ASW i.e., 100%, 90%, 80%, 70%, 60%, and 50% which correspond to 3%, 2.7%, 2.4%, 2.1%, 1.8%, and 1.5% salinities respectively. Data represent means ± SEM for 3 independent 15 mL batch cultures (n=3). Asterisks indicate significant differences between the methods tested using two-way ANOVA (Tukey’s test); * p<0.05, ** p<0.01, and *** p<0.001.

### *R. sulfidophilum* shows comparable growth, nutrient, and amino acid profile in 40% ASW-based heterotrophic culture medium (1.2% salinity) without additional contamination risks

To evaluate the impact of lower salinities (50%, 40%, and 30% ASW corresponding to 1.5%, 1.2%, and 0.9% salinities respectively) on the growth of *R. sulfidophilum* on a larger scale, it was cultured in ASW supplemented with 0.1% yeast extract and 0.5% peptone in 10 L bottles (Fig. 2a). The dry cell yields tended to be higher in 50% and 40% ASW, but lower in 30% ASW compared to 100% ASW (Fig. 2b). Thus, 40% ASW or 1.2% salinity was used as the low salinity medium for testing growth, nitrogen : phosphorus : potassium (N : P : K) and the amino acid profile in 10 L scale to evaluate the implications of this reduction on its use as a plant N fertilizer. The initial and final OD_660_ (Fig. 2c) and the initial and final dry cell yield (Fig. 2d) were not different between the 100% and the 40% ASW treatments, but tended to be higher in the lower salinity medium (40% ASW). Further, there were no additional risks of contamination caused by lowering the salinity to 1.2% (Fig. 2e). The biomass obtained from the respective treatments was lysed and dried (<u>P</u>rocessed <u>B</u>iomass or PB hereafter) for further analysis. The N : P : K of PB from 40% ASW treatment was 11.9 : 2.94 : 0.57, comparable to previously reported values in MB (11.0 : 2.95 : 0.51) (18), except for N which tended to be higher. The total amino acid composition between the PB from 100% and 40% ASW was comparable, except for glutamate and glutamine, which were higher in 40% ASW (Table 2). The composition of free amino acids, on the other hand, was marginally different, with higher lysine, glutamate, and aspartate; and lower arginine in PB from 40% ASW (Table 2).

**Fig. 2.**
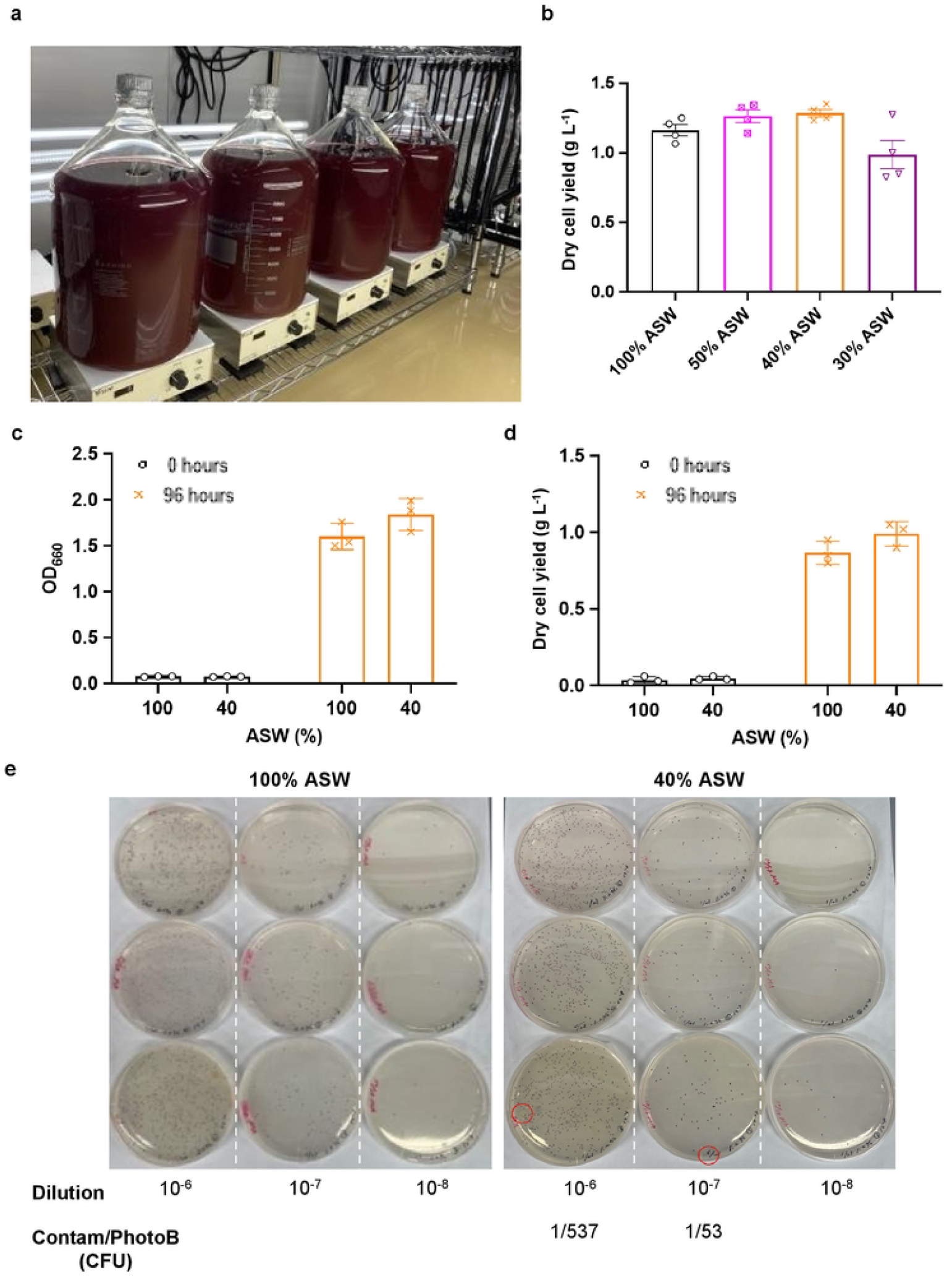
Dry cell yield and contamination status in ASW-based *R. sulfidophilum* cultures at a reduced salinity of 1.2%; 100% and 40% correspond to 3% and 1.2% salinity respectively. **a**, Ten-liter culture set-up and **b**, dry cell yield (g L^-1^) of *R. sulfidophilum* in ASW supplemented with 0.1% yeast extract and 0.5% peptone at 100%, 50%, 40%, and 30% ASW, corresponding to 3%, 1.5%, 1.2%, and 0.9% salinities respectively. **c**, Initial and final OD at 660 nm and **d**, the dry cell yield (g L^-1^) were obtained for 100% and 40% ASW treatments. **e**, Contamination status at the end of the culture period was obtained for 10^−6^ to 10^−8^ serial dilutions of 100% (*left panel*) and 40% (*right panel*) ASW treatments. Data represent means ± SEM for n=3 or 4. Significant differences between the treatments tested were not observed. Red circles indicate the contaminating colonies.

**Table 2.** Free and total amino acid profile expressed in weight % of lysed and dried biomass (Processed Biomass or PB) from *R. sulfidophilum* cultured in 100% and 40% ASW. Data are shown for the combined PB from 3 independent batch cultures. n.d. indicates no data due to degradation by HCl.

| Amino acid | Total amino acid content (weight %) |  | Free amino acid content (weight %) |  |
| --- | --- | --- | --- | --- |
|  | 100% ASW | 40% ASW | 100% ASW | 40% ASW |
| Arg | 7.62 | 7.42 | 37.56 | 30.28 |
| Asp + Asn | 10.65 | 10.47 | 14.11 | 15.77 |
| Glu + Gln | 13.73 | 14.54 | 12.43 | 14.68 |
| Lys | 5.40 | 5.54 | 16.33 | 20.42 |
| Ser | 3.54 | 3.41 | 0.29 | 0.12 |
| Thr | 4.77 | 4.76 | 0.50 | 0.45 |
| His | 1.87 | 1.84 | 0.00 | 0.00 |
| Met | 3.11 | 3.11 | 0.64 | 0.62 |
| Ala | 9.20 | 9.05 | 1.56 | 1.66 |
| Val | 6.84 | 6.80 | 1.74 | 1.33 |
| Gly | 5.86 | 5.82 | 3.09 | 3.02 |
| Ile | 4.83 | 4.77 | 1.93 | 2.14 |
| Leu | 9.43 | 9.32 | 3.96 | 3.37 |
| Phe | 5.36 | 5.36 | 3.05 | 3.18 |
| Pro | 4.25 | 4.19 | 1.00 | 1.24 |
| Trp | n.d. | n.d. | 0.41 | 0.43 |
| Tyr | 3.53 | 3.60 | 1.21 | 1.10 |

### 40% ASW can replace 100% ASW for the autotrophic growth of *R. sulfidophilum*

To test if 40% ASW could replace the 100% ASW in autotrophic conditions, *R. sulfidophilum* was cultured in 100% and 40% ASW supplemented with 10 mM sodium thiosulfate pentahydrate and a CO_2_/N_2_ (7:3) gas mixture for 4 days in 2 L jar fermenters (Fig. 3a). Cells grew similarly (based on OD_660_) between the two treatments (Fig. 3b), and a comparable final dry cell yield of ∼0.4 g L^-1^ was obtained for the two treatments (Fig. 3c). However, 40% ASW i.e., the 1.2% salinity condition tended to display higher initial growth than 100% ASW i.e., the 3% salinity condition (Fig. 3b and 3c). Specifically, the growth rate during the first day of culture was higher at 1.2% salinity, while the growth rates on day 2 and day 3 remained higher at 3% salinity (Fig. 3d). Similar trends were also observed with the total organic carbon (TOC) and the total nitrogen (TN) assimilated by cells which were obtained from the sole C and N sources, CO_2_ and N_2_, respectively. TOC of cells in 1.2% salinity was higher during initial growth (Fig. 3e), consistent with cell growth (Fig. 3b and 3c). However, the TN remained higher in cells throughout the culture in 1.2% salinity compared to 3% salinity (Fig. 3f).

**Fig. 3.**
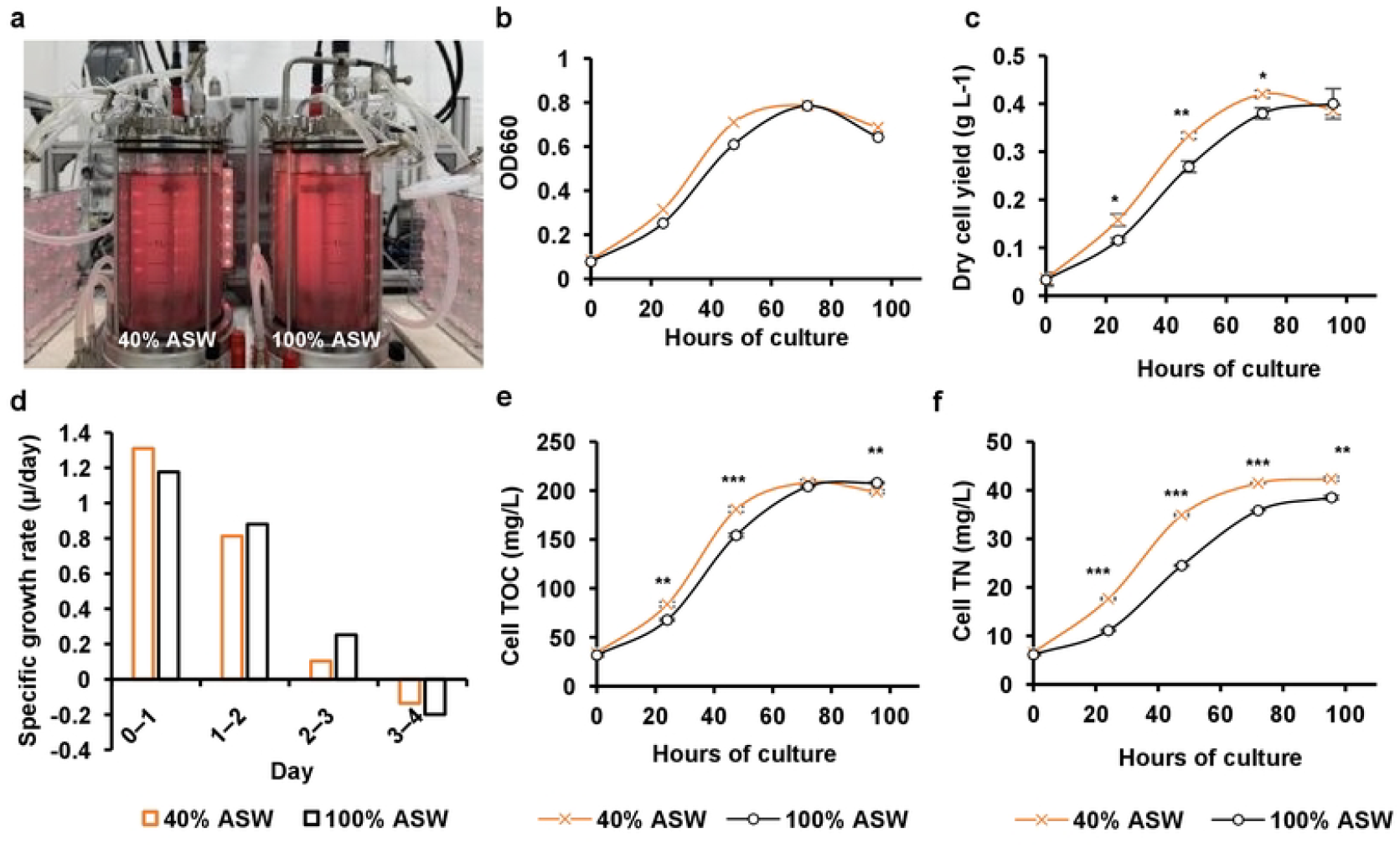
Cell growth, dry cell yield, total organic carbon (TOC), and total nitrogen (TN) in autotrophic cultures of *R. sulfidophilum* in artificial seawater (ASW) based growth medium of different salinities, wherein 100% corresponds to 3% salinity and 40% corresponds to 1.2% salinity. **a**, Two-liter fermenter set-up for the culture of *R. sulfidophilum* in 100% or 40% ASW supplemented with 10 mM sodium thiosulfate pentahydrate and CO_2_/N_2_ (7:3) gas mixture. **b**, Optical density (OD) at 660 nm, **c**, dry cell yield (g L^-1^), and **d**, OD-based specific growth rates (*μ*) were obtained for every day of the culture. The total C and N in the cell fractions are shown as **e**, cell TOC (mg L^-1^), and **f**, cell TN (mg L^-1^) respectively. Data represent means ± SEM (n=3). Asterisks indicate significant differences between the methods tested using the student’s T-test statistic; * p<0.05, ** p<0.01, and *** p<0.001.

## Discussion

We evaluated the performance of *R. sulfidophilum* in a low-salinity growth medium as a cost-effective alternative to MB and confirmed its effectiveness as a plant N fertilizer. Lowering the salinity to 1.2% did not adversely affect cell growth and the N, P, K content, but tended to improve dry cell yield (Fig. 2d and 3c) and N content, and alter the amino acid composition (Table 2). While one of the advantages of using natural seawater (salinity of ∼3% or above) as a culture medium is its low risk of contamination, decreasing the salinity of seawater-based medium to 1.2% did not increase the contamination events during cultivation in 10 L-scale (Fig. 2e). Thus, the culture medium for large scale cultivation of *R. sulfidophilum* was comprehensively improved without any trade-offs.

The trend of higher growth (Fig. 1a) and dry cell yield in culture medium with lower salinities (Fig. 1b and 1c) is consistent with a previous study, wherein decreased PHA accumulation in cells was also observed with a decrease in salinity from 4.5% to 1.5% (19). Acetyl-CoA is the precursor for PHA biosynthesis in *R. sulfidophilum* (9,15) and also a common metabolic intermediate of the C metabolism feeding into the TCA cycle, which provides C skeletons necessary for N metabolism including the biosynthesis of amino acids (20). Hence, a decrease in C flux towards PHA biosynthesis could increase the flux towards the TCA cycle, facilitating improved growth and/or N assimilation in the lower salinity growth medium. This was also observed in the current study wherein cells when grown autotrophically, tended to assimilate more N in the 1.2% salinity growth medium (Fig. 3f). A similar tendency may also be expected in heterotrophic cultivation leading to marginally higher N content (11.9%) in PB at 1.2% salinity. However, due to the difference in the modes of metabolism, further study would be necessary to verify the mechanism of a higher N utilization and N content in a 1.2% salinity medium under respective growth conditions.

Furthermore, PB from a 1.2% salinity growth medium possessed higher free lysine and glutamate (Table 2), which are reported to play a role in the induction of stress responses in tomato (21) and rice (22) respectively. Free aspartic acid, exogenous application of which has been reported to mitigate the adverse effects of salt stress in wheat (23), also tended to be higher in a 1.2% salinity growth medium. However, free arginine, which plays a minimal role in stress resilience but is a good N source (24,25) was lower in PB from 1.2% salinity growth medium. Although these findings point towards the possible function of PB in enhancing stress resilience in plants, further experiments would be required to confirm its simultaneous use as a fertilizer and a biostimulant.

## Methods

### Cultivation of *R. sulfidophilum*

*R. sulfidophilum* (DSM 1374; W4, LMG 5202) (ATCC) was grown in various media with different salt compositions as detailed below. The cultures were maintained at room temperature and 80 W/m^2^ (Y104-R660/W40/IR73-31W-EI0U1LW-010, YUMEX Solutions) with a starting OD_660_ of 0.1.

#### Seawater based medium

Media ranging from 50% (15 g L^-1^) to 100% (30 g L^-1^) ASW (Marine standard, Nihon Kaisui Co. Ltd, Tokyo, Japan) in 10% increments were supplemented with either 0.1% (w/v) yeast extract and 0.5% (w/v) peptone or one mM sodium acetate and 2.5 mM sodium thiosulfate, and used to culture *R. sulfidophilum* to obtain the growth curve and cell dry weights. NSW harvested from Maizuru Bay was passed through 5 μm and 0.5 μm polypropylene wound cartridge filters (Advantec, Tokyo, Japan) into 6 UV sterilizers (UV40W, Kankyo Technos Co. Ltd., Wakayama, Japan) connected in tandem, and circulated through this system periodically to prevent contamination during storage. The cultures in NSW supplemented with 0.1% (w/v) yeast extract and 0.5% (w/v) peptone were used for cell yield and cost analysis comparisons with MB and ASW-based medium. All media were sterilized in an autoclave at 121°C at 15 psi for 30 minutes (10 L scale). Cultures in 100% and 40% ASW supplemented with 0.1% (w/v) yeast extract and 0.5% (w/v) peptone in a 10 L culture scale were used for nutrient (N, P, K) and amino acid analysis.

### Growth analysis and yield of dry cell weight

The OD_660_ readings were taken every 24 hours to plot growth curves for three independent biological replicates under each salinity condition. A fixed volume of 7-day-old (Fig. 1) or 4-day-old cell cultures (Fig. 2 and Fig. 3) were centrifuged at 9000 x*g* for 10 minutes, washed with 1 ml sterile water, transferred to pre-weighed empty 1.5 ml centrifuge tubes, and centrifuged again at 9000 x*g* for 20 minutes. The pellets were lyophilized to determine cell dry weights (g), which were then converted to yield (g L^-1^) based on the volume of the culture used for centrifugation.

### Contamination check, harvest, lysis and drying

Contamination was checked by plating 100 μl of 10^−6^, 10^−7^ and10^−8^ serial dilutions of 4-day-old cultures on marine agar [Marine Broth (Merck, Millipore Sigma, Massachusetts, United States) with 1.5% Agar], and incubating at 30°C for 3 days under far-red light (25 W m^-2^). Colony forming units (CFU) of *R. sulfidophilum* and contaminants were counted on plates showing contamination.

Cells from 4-day-old 10 L cultures were first flocculated in 7.5 mg L^-1^ chitosan 100 (Fujifilm Wako, Osaka, Japan) in 0.01% acetic acid (Fujifilm Wako) for 2–3 days, which were then harvested by centrifugation (14,000 x*g*, 10 minutes), and resuspended in pure water to remove the salt. For cells cultured in marine broth, 12 mg L^-1^ of chitosan 100 in 0.01% acetic acid was used for flocculation. Washed cells were then collected by centrifugation (14,000 x*g*, 60 minutes) and stored at –80°C until further use. Cell pellets from 3 independent culture bottles in each salinity condition were combined, chemically lysed, air-dried, and powdered for nutrient and amino acid analysis.

### Nutrient and amino acid analysis

Lysed and dried bacterial biomass was analyzed for N and C content by dry combustion. Phosphate (P_2_O_5_) and potassium oxide (K_2_O) were extracted through nitric acid decomposition and analyzed using absorptiometry and atomic absorption spectroscopy, respectively, at Vegetech Co., Ltd. Physical and Chemical Analysis Center, Kanagawa, Japan. Values are expressed as % dry weight.

The amount of free (not bound in a protein) and total (proteinogenic + free) amino acids (except for cysteine and tryptophan) in the lysed and dried bacterial biomass were quantified with an amino acid analyzer (L-8900, Hitachi) using the post-column ninhydrin derivatization of amino acids. For the quantification of free amino acids, 5 mg of the air-dried sample was subjected to sonication in 200 μL of 0.2 M perchloric acid for ∼2 minutes and incubated at 0°C for 30 min. Following centrifugation at 12,000 x*g* for 5 minutes, the supernatant was diluted 3.3-fold with lithium citrate buffer (pH 2.2) (123-02505, Fujifilm Wako Pure Chemical Corporation), and 90 μL was used for the analysis. For the quantification of total amino acids (18 of the 20 proteinogenic amino acids, excluding cysteine and tryptophan), 1 mg of the lyophilized sample was subject to hydrolysis in 100 μL of 6 M hydrochloric acid at 110°C for 20 hours. The volume of the hydrolysate was adjusted to 100 μL with ultra-pure water and filtered through a 0.45 μm filter. The filtrate was diluted 20-fold with the lithium citrate buffer and 15 μL was used for the analysis.

### Autotrophic cultivation and Total organic Carbon (TOC)/ Total Nitrogen (TN) analysis

*R. sulfidophilum* was autotrophically cultured in 100% and 40% ASW supplemented with 10 mM sodium thiosulfate pentahydrate and a CO_2_/N_2_ (7:3) gas mixture in two independent 2 L jar fermenters (BEM, Tokyo, Japan) at 27°C for 4 days under far-red light (25 W m^-2^). The culture medium was bubbled with N_2_ for 15 minutes at 0.5 L min^-1^ to purge the dissolved oxygen. The gas mixture was then bubbled at 0.5 L min^-1^ through the culture medium for 30 minutes, and the pH of the culture was maintained at 7.5 using 1N NaOH. The gas exhaust line was then sealed to establish a closed system. Following the bubbling, sodium thiosulfate pentahydrate and the seed culture of *R. sulfidophilum* cultured in M6 minimal medium, as previously reported (10), were inoculated into the culture medium to a final OD_660_ of 0.1. Samples were collected at the start of the culture (0 hours) and 24 hours, 47.5 hours, 72 hours, and 95.5 hours of culture to evaluate growth (OD_660_), dry cell yield, total organic carbon (TOC), and total nitrogen (TN) content. TOC and TN were obtained for ∼43 ml of culture and the culture medium, and the samples for culture medium at corresponding time points were prepared by centrifugation of cultures at 9000 x*g* for 10 minutes. TOC and TN were analyzed by TOC-L and TNM-L (Shimadzu, Kyoto, Japan). TOC was obtained by subtracting the inorganic C from the total C present in the culture or the culture medium (supernatant) at a given time point. The amounts of C and N fixed by the bacteria were calculated by subtracting the amounts of TOC or TN in the culture medium (supernatant) from the TOC or TN in the culture at each corresponding time point. The cells were harvested and processed for the N, P, K analysis as mentioned in the earlier sections.

## Acknowledgments

We would like to thank Dr. Nehlah Rosli (Symbiobe Inc.) for her assistance with preculture preparation used in the autotrophic cultivation of *R. sulfidophilum*. This work was supported by Japan Science and Technology, COI-Next (Grant Number JPMJPF2114). The funder played no role in the study design, data collection, analysis, interpretation, or writing of this manuscript.

## Data availability

All data generated or analyzed during this study are included in this published article and its supplementary information files.

## Author contributions

KN conceived the original research idea; SRM-Y, DDH, MS, SK, and GL designed the experiments. SRM-Y, DDH, MS, YK, and AY performed the experiments. HM, SK, MO, and GL obtained and analyzed the amino acid data. SRM-Y analyzed all data and wrote the manuscript with contributions from all authors. KN supervised the study. All authors read and approved the final manuscript.

